# Regulation of immune signal integration and memory by inflammation-induced chromosome conformation

**DOI:** 10.1101/2024.02.29.582872

**Authors:** Bence Daniel, Andy Y. Chen, Katalin Sandor, Wenxi Zhang, Zhuang Miao, Caleb A. Lareau, Kathryn E. Yost, Howard Y. Chang, Ansuman T. Satpathy

## Abstract

3-dimensional (3D) genome conformation is central to gene expression regulation, yet our understanding of its contribution to rapid transcriptional responses, signal integration, and memory in immune cells is limited. Here, we study the molecular regulation of the inflammatory response in primary macrophages using integrated transcriptomic, epigenomic, and chromosome conformation data, including base pair-resolution Micro-Capture C. We demonstrate that interleukin-4 (IL-4) primes the inflammatory response in macrophages by stably rewiring 3D genome conformation, juxtaposing endotoxin-, interferon-gamma-, and dexamethasone-responsive enhancers in close proximity to their cognate gene promoters. CRISPR-based perturbations of enhancer-promoter contacts or CCCTC-binding factor (CTCF) boundary elements demonstrated that IL-4-driven conformation changes are indispensable for enhanced and synergistic endotoxin-induced transcriptional responses, as well as transcriptional memory following stimulus removal. Moreover, transcriptional memory mediated by changes in chromosome conformation often occurred in the absence of changes in chromatin accessibility or histone modifications. Collectively, these findings demonstrate that rapid and memory transcriptional responses to immunological stimuli are encoded in the 3D genome.

## Introduction

Innate immune cells represent the first line of defense against pathogens and evolved to rapidly interpret and respond to changes in their environment. Macrophages (MFs) are one of the most plastic cellular components of the innate immune system and possess the ability to alter their phenotype, for example, by integrating multiple co-existing signals to produce the appropriate inflammatory response, or by establishing immune memory following pathogen clearance^1–3^. MF plasticity can be programmed by multiple molecular inputs: 1) cell surface receptors that recognize distinct pathogen-associated molecular patterns; 2) regulation of signal-dependent transcription factors (TFs) downstream of receptor signaling, and 3) a highly plastic and permissive epigenetic landscape that encodes transcriptional responses to receptor signaling and TF pathways^2^. In the first few hours of inflammation, MFs undergo polarization and change their phenotype by regulating the expression of hundreds of genes^4–8^. This rapid transcriptional response is the consequence of a primed epigenetic state, in which, accessible, gene-distant regulatory elements (enhancers) are brought in close proximity to gene promoters to induce gene expression^8–12^. Therefore, the 3-dimensional (3D) conformation of the genome is presumed to be a critical part of regulating the inflammatory response; however, the relationship between genome architecture and its functional impact on the scale and timing of the inflammatory response, resolution of inflammation, and memory, is understudied.

The genome is organized into large-scale 3D structures, such as compartments and topologically associating domains (TADs – demarcated by the insulator binding TF, CTCF), that separate transcriptionally/epigenetically active (Compartment A) and repressed (Compartment B) genomic regions, and genes with their enhancer elements, respectively^13–15^. In MFs, our current understanding supports the notion that compartments and TADs are largely inert to external signals; however, Cohesin and some CTCF boundaries seem to be crucial for signal-driven, rapid transcriptional responses in immune cells, including MFs^19–21^. At the level of enhancer-promoter communication, chromatin interactions are considered to be largely preformed (primed) to support quick and robust transcriptional responses to inflammation in innate immune cells^22^; however, *de novo* loop formation has been reported in the context of bacterial infection, inflammation, and interferon-driven MF responses^23–25^. These findings indicate that chromatin conformation can exhibit flexibility in a changing microenvironment, such as during an innate immune response; although we lack knowledge – mainly due to technological limitations (i.e., resolution to identify enhancer-promoter loops) – on whether inflammation-induced genome conformation can regulate cellular plasticity and memory formation^26^.

Traditional 3C-based approaches, such as Hi-C, can map large-scale architectural features of the genome with high confidence (TADs and compartments); however, identification of specific gene-regulatory chromatin loops (i.e., enhancer-promoter contacts) with high-resolution is more challenging^16^. Recent studies using micrococcal nuclease-based genome fragmentation following proximity ligation, such as Micro-Capture C (MCC), enabled the generation of base-pair resolution contact maps of specific genomic regions^17,18^. Exploiting these technological advances, here, we investigate the roles of 3D genome conformation in the setting of IL-4-induced alternative macrophage polarization, inflammatory signal integration, and short-term memory formation. IL-4 is a T helper 2 type cytokine that is critical for defense against multicellular parasite infections, drives alternative (M2) MF polarization, can imprint a stable 1-dimensional (1D) epigenomic program (i.e., histone modifications, TF binding, and chromatin accessibility) during MF polarization with lasting effects to alter the subsequent inflammatory response to endotoxin and interferons^3^. Importantly, the interaction between IL-4 and endotoxin can have detrimental, cytokine storm-like effects in the *in vivo* murine model of airway inflammation and asthma that was suggested to be the result of 1D epigenomic remodeling^3, 7, 8, 11, 30^.

In this work, global genome conformation analysis with Hi-C demonstrated that IL-4 creates ∼7,000 chromatin loops during MF polarization (in 24-hours), of which, ∼50% show short-term memory (stability) after the removal of the cytokine. Of these genome conformational changes, we find thousands of IL-4-induced chromatin loops that do not associate with 1D epigenomic-, or transcriptional-changes, some that occur in the proximity of inflammatory genes, such as *Il6*. To study the functional role of IL-4-induced chromatin loops, we establish two treatment paradigms: 1) to study signal integration (two signals are present at the same time) between IL-4 and a secondary, unrelated signal (Lipopolysaccharide – LPS; Interferon-gamma – IFNG; and Dexamethasone – DEX), and 2) to study short-term transcriptional memory formation that is induced by IL-4 and affects the outcome of MF responses to the above-mentioned secondary signals. We report vastly reprogrammed transcriptional responses following IL-4 treatment that associate with a remodeled 3D genome conformation of specific gene loci. Using high-resolution MCC analysis to infer promoter interaction profiles of multiple genes, we report that IL-4 primes 3D genome conformation by juxtaposing LPS-, IFNG-, and DEX-activated enhancers to their cognate gene promoters. CRISPR-deletion of the anchor points of IL-4-induced chromatin conformation (i.e., enhancers and CTCF-bound insulators) proved that these *de novo* genome architectural features are required for enhanced and synergistic LPS-driven transcriptional responses, and can provide short-term transcriptional memory. Collectively, our work provides evidence that MF priming occurs in the 3D genome, the conformation of the DNA can integrate inflammatory signals, and cellular memory can be imprinted in the 3D architecture of DNA.

### Macrophage polarization stably remodels 3D genome conformation

MFs undergo alternative polarization in the presence of IL-4, a cytokine with stable, long-term 1D epigenomic effects that change MF plasticity to unrelated environmental signals via transcriptional memory^7, 8, 27–30^. Thus, we hypothesized that IL-4 remodels the 3D genome conformation of MFs, and this genome conformational change affects MF plasticity. To better understand whether IL-4 impacts genome conformation, we set up a MF priming model in which MFs are exposed to IL-4 for 24-hours (IL-4), followed by washout, and a 24-hour resting period (IL-4-primed – pIL-4). Using these three conditions, we performed Hi-C, RNA-seq, ATAC-seq and ChIP-seq for three histone marks that associate with active chromatin (H3K27ac, H3K4me1, and H3K4me3) to gather information about IL-4 initiated genome conformational, 1D epigenomic, and transcriptomic changes during priming. We performed differential loop calling from Hi-C data (10kb resolution, Log_2_FC>0.5, p-value<0.1), and identified a total of 10,438 differential loops among the three conditions, that included 6,923 IL-4-induced and 3,515 IL-4-repressed loops, respectively (**Figure 1A; Figure S1A)**. We further analyzed the IL-4 induced loops, and identified three IL-4-induced loop patterns by k means clustering: 1) Transient – lost after IL-4 removal (n=1,776; e.g., *Arg1, Itgax*), 2) Primed – gained mainly after IL-4 removal (n=1,720; e.g., *Ccl2, Il6*), and 3) Memory – retained after IL-4 removal (n=3,427; e.g., *F7, F10*) (**Figure 1A)**. These results report IL-4-induced chromatin loops that exhibit stability in the MF priming model.

**Figure 1.**
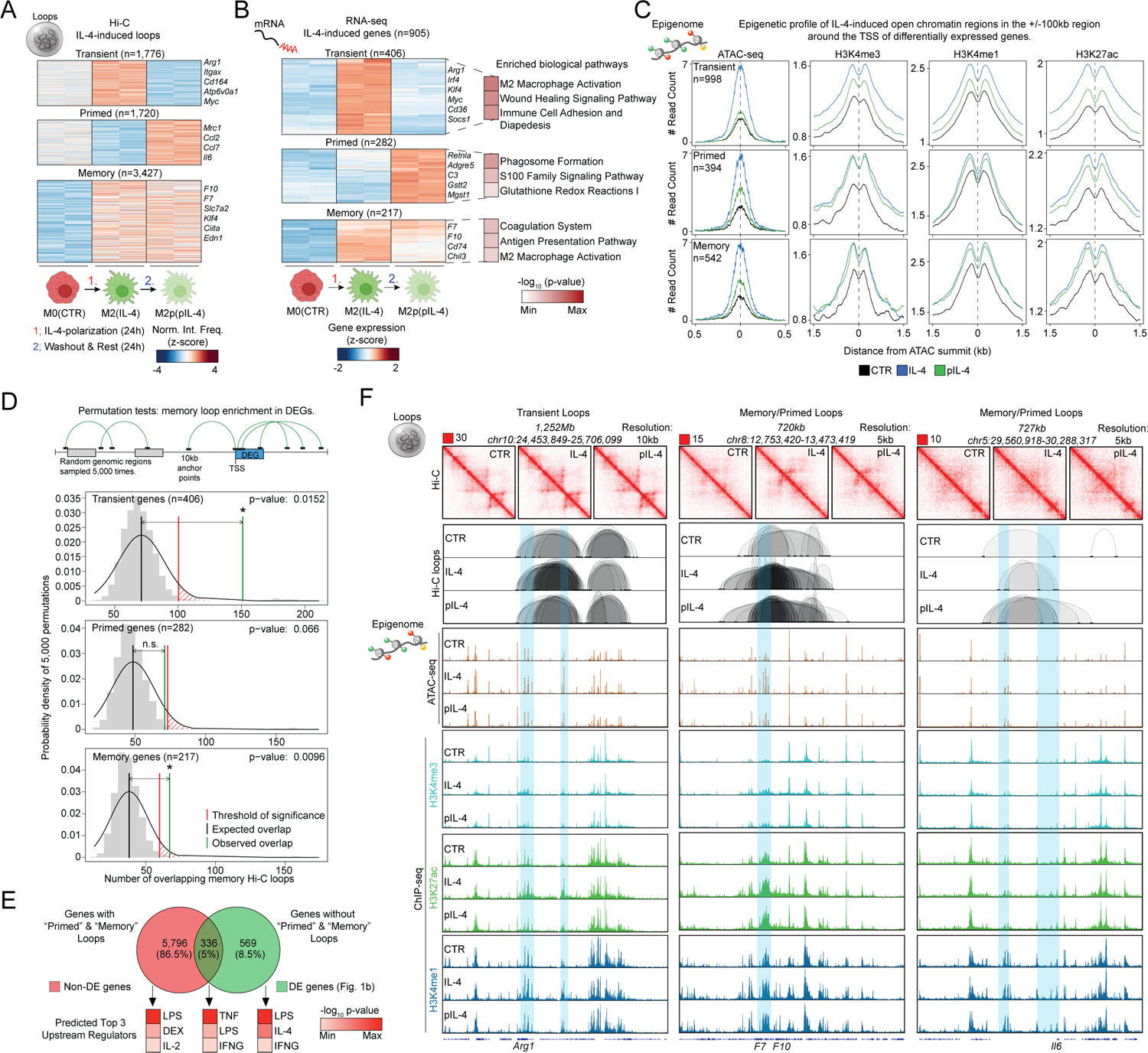
IL-4 remodels chromosome conformation with memory potential. **(A)** Heatmap representation of differential chromatin loops detected by Hi-C. **(B)** Heatmap representation of differentially expressed genes (DEGs) detected by RNA-seq (left). Small heatmaps represent the results of Ingenuity pathway analysis for each RNA-seq gene cluster, and the top 3 biological terms are shown (right). **(C)** Histograms represent the average ATAC-seq and ChIP-seq signal at IL-4 induced open chromatin regions that are in the proximity of DEGs from panel b. **(D)** Scheme of the permutation test (top). Permutation test results of the enrichment of memory loops that overlap with DEGs or random genomic regions that are sampled 5,000 times. **(E)** Venn diagram represents the overlap between DEGs and non-DEGs that associate or do associate with Memory loops. Ingenuity upstream regulator analysis of the gene sets in the Venn diagram with the top 3 cytokine and chemical drug upstream regulators. **(F)** Genome browser views of the indicated datasets in the *Arg1, F7/F10*, and *Il6* genomic loci.

Next, we performed differential gene expression analysis from RNA-seq data (FDR<0.05, Log_2_FC>0.5) and using k means clustering, we identified gene groups with similar gene expression patterns to the above-described chromatin loop patterns, featuring IL-4-induced Transient (n=406), Primed (n=282), and Memory (n=217) gene expression patterns (**Figure 1B**, **Table S1**). Biological pathway analysis (Ingenuity Pathway Analyzer) showed that Transient genes were enriched for Alternative or M2 MF activation-related terms, Primed genes were enriched for inflammation-associated terms, such as Phagosome formation and S100 Signaling Pathway, while Memory genes were enriched for Coagulation-, Antigen presentation-, and M2 MF activation-related terms (**Figure 1B)**. IL-4 repressed genes (n=764) were largely enriched for inflammation-related terms, such as Toll-like receptor signaling and M1 MF activation **(Figure S1B, Table S2)**^8^.

To characterize the 1D epigenomic changes that occur during IL-4-priming, we performed differential open chromatin region (OCR; p-value<0.05, Log_2_FC>0.5) and differential histone modification enrichment analyses (H3K27ac H3K4me3, and H3K4me1; p-value<0.05, Log_2_FC>0.5), and found similar 1D epigenomic patterns to the ones that we described above for gene expression (k-means clustering; **Figures S1C and S1D**). To investigate the epigenetic state of the above-described gene groups, we linked IL-4-induced OCRs (n=8,269) to these genes (+/- 100kb around the transcription start sites - TSSs; scheme - **Figure S1E**), resulting in IL-4-induced OCR sets of Transient (n=998), Primed (n=394), and Memory (n=542) genes. We observed that IL-4-induced chromatin accessibility was largely transient in all three gene groups, and stability, or memory was mainly detected when histone modifications were analyzed, except for Transient genes, where we observed that all three histone marks exhibited transient pattern (**Figure 1C; Figures S1C and S1D)**. Of note, among the 1D epigenetic features we studied, H3K4me1 exhibited by far the most stability after washout, retaining ∼30% (n=1,217) of IL-4-induced signal in the genome compared to the other two histone marks (H3K4me3, ∼10%, n=174; H3K27ac, ∼19%, n=794) or open chromatin (∼10%, n=859; **Figures S1C and S1D**). These results indicate that IL-4-induced genome conformation exhibits stability similarly to certain histone modifications and might underlie memory function.

### IL-4-driven 3D genome reprogramming affects inflammatory gene loci in the absence of transcriptional and 1D epigenomic changes

To couple 3D genome conformational changes to gene expression, we performed permutation tests to evaluate whether memory loops from Hi-C are significantly enriched in the gene bodies of genes that exhibit Transient, Primed, and Memory gene expression patterns over random genomic regions sampled 5,000 times. We found that Memory loops were enriched in the vicinity of Transient and Memory genes, but not in Primed genes (**Figure 1D**). Next, we assessed how many genes in the genome overlap with Memory loops, Primed loops, or both. We found that 336 DEGs (from Figure 1B) showed overlap with Memory/Primed loops (e.g., *F7* and *F10*), while 569 did not. Additionally, we found 5,796 non-DEGs with overlapping Memory/Primed loops, including pro-inflammatory genes, such as *Il6* (**Figures 1E and 1F; Table S3**). We performed upstream regulator analysis of DEGs without Memory/Primed loops, DEGs with Memory/Primed loops, and non-DEGs with Memory/Primed loops, and found that lipopolysaccharide (LPS), dexamethasone (DEX), tumor necrosis factor (TNF), interferon gamma (IFNG) and interleukin-4 (IL-4) are the top predicted upstream regulators of these gene sets with LPS, DEX and IL-2 being predicted for the 5,796 non-DEGs with Memory loops (**Figure 1E**). Next, we examined how many of the non-DEGs associate with IL-4-induced OCRs, and we found that ∼60% had IL-4-induced OCRs in their vicinity (+/- 100kb around the TSSs; OCR - n=3,961) with stable histone mark changes, indicating that these genes undergo transcriptionally silent, 3D epigenomic reprogramming, whereas the remaining 40% undergo transcriptionally silent, and largely 1D epigenetically silent - according to the epigenetic features we studied – (OCR – n=3,696), 3D genome reorganization (**Figure S1F)**. These results suggest the uncoupling of chromatin accessibility, histone modifications, and 3D genome-reprograming from gene expression as part of MF priming/training, and raises the notion that IL-4-induced genome conformation has functional importance, perhaps when these primed/trained MFs encounter subsequent environmental signals (e.g., LPS, DEX, IFNG), which is, to date, have been mainly linked to epigenetic priming/training^7, 8, 11, 30^.

### IL-4-driven 3D genome conformation associates with enhanced response to endotoxin and dexamethasone

To better understand the roles of IL-4-induced 3D genome conformation changes in MF responses to environmental challenges, we set up two model systems to study how MFs integrated two environmental signals at the same time (Signal integration), and to assess whether IL-4 can imprint short-term transcriptional memory via genome conformation (Memory). We induced M2 MF polarization (24-hours IL-4), and then either: 1) immediately treated MFs for three-hours with Lipopolysaccharide (LPS), Interferon gamma (IFNG), or Dexamethasone (DEX) (referred to as secondary stimulants) in the presence of IL-4 (Signal Integration), or 2) washed out IL-4, rested the cells for 24-hours and treated with the secondary stimulants (Short-term memory) (**Figure 2A**). We performed bulk RNA-seq in these settings in wild type and *Stat6^-/-^* mice. Differential gene expression analysis (FDR<0.05, Log_2_FC>0.5) revealed hundreds of genes with enhanced response to all three secondary stimulants following IL-4 priming in both the Signal integration and Short-term transcriptional memory formation models. Namely, we found 849 genes with enhanced LPS response (k means clustering; Cluster 1 – strong synergistic response in both models; Cluster 2 – enhanced/synergistic response in both models, no IL-4-induced transcription; Cluster 3 – strong synergistic response in the model of memory, no IL-4-induced transcription; **Table S4**), 536 genes with enhanced DEX response (k means clustering; Cluster 1 - strong synergistic response in the model of memory, no IL-4-induced transcription; Cluster 2 - enhanced/synergistic response in both models; Cluster 3 – strong synergistic response in signal integration; **Table S5**), and 215 genes with enhanced IFNG response (k means clustering; Cluster 1 – enhanced response in both models, no IL-4-induced transcription; Cluster 2 – enhanced response in signal integration with partial memory response; **Table S6**). Genes with an enhanced response after LPS treatment showed enrichment for cell adhesion (Cluster 1 - e.g., *Rhob, Rhoc*), classical MF activation (Cluster 2-e.g., *Cd86, Il6, Slc7a2*), and cholesterol metabolic pathways (Cluster 3 - e.g., *Cyp51a1, Hmgcr*); genes with enhanced DEX response showed enrichment for MF-related inflammatory functions and alternative M2 MF activation (Clusters 2 and 3 - e.g., *Klf4, Mrc1, Fabp4*); and genes with enhanced IFNG response were enriched for inflammation-related signaling pathways, including cytokine storm signaling (Cluster2 - e.g., *Ccl2, Ciita, Ccl11*; **Figure 2B, C, and Figure S2A**).

**Figure 2.**
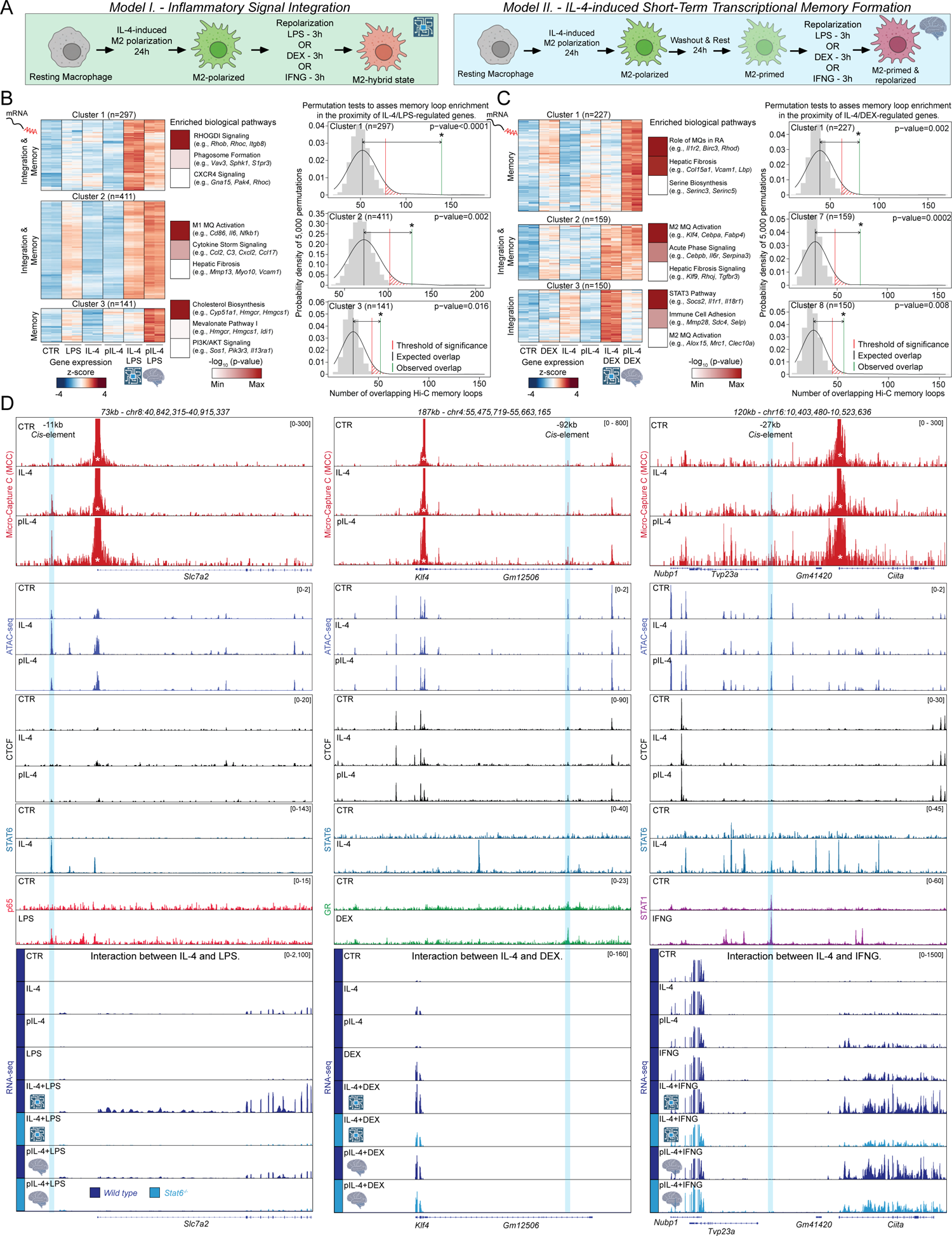
IL-4-induced chromosome conformation juxtaposes signal-responsive enhancers and their cognate promoters. **(A)** Schematics of the two model systems used to study Inflammatory signal integration (Model I.) and Short-term Memory formation (Model II.). **(B)** Heatmap representation of differentially expressed genes (DEGs) detected by RNA-seq (left). Small heatmaps represent the results of Ingenuity pathway analysis for each RNA-seq gene cluster, and the top 3 biological terms are shown (middle). Permutation test results that assess the enrichment of Memory loops in the different groups of DEGs in the IL-4 – LPS treatment paradigm. **(C)** Same as B, for the IL-4 – DEX treatment paradigm. **(D)** Genome browser views in three different genomic loci that show how IL-4 priming establish specific looping interactions between gene promoters and *cis*-elements that are bound by transcription factors activated by secondary stimulants and associate with enhanced transcriptional responses.

We performed permutation tests to assess whether Memory and/or Primed loops detected by Hi-C were enriched in the gene bodies of genes exhibiting enhanced response to LPS, DEX and IFNG, and found that all three gene clusters in both the IL-4 – LPS (permutation tests: Cluster 1 p<0.0001; Cluster 2 p<0.002; Cluster 3 p<0.016) and IL-4 – DEX (permutation tests: Cluster 1 p<0.002; Cluster 2 p<0.0002; Cluster 3 p<0.008) treatment settings showed enrichment for Memory/Primed loops (**Figure 2B and C**). We did not find a significant enrichment of these loops in genes that show enhanced IFNG response after IL-4 treatment, likely due to the fact that there are fewer genes that exhibit signal integration and/or short-term transcriptional memory in this system compared to LPS and DEX (gene counts - IFNG: 215, DEX: 536, LPS: 849; **Figure S2A**). We also did not observe Memory and/or Primed loop enrichment in genes that show dampened response to LPS, DEX and IFNG, except for one gene cluster which showed reduced LPS response (Cluster 5; **Figure S2B, Table S7**). These results suggest that IL-4-induced Memory loops are enriched near genes that show enhanced gene expression profiles after IL-4 priming and nominated genes for further analysis to better understand the roles of Memory loops in signal integration and short-term transcriptional memory formation.

### IL-4 induced memory loops bridge signal-responsive enhancers and promoters

To precisely pinpoint the anchor points of IL-4-induced memory loops, we performed Micro-Capture C (MCC) experiments to assess the inflammatory response of primary MFs. This method can generate base-pair resolution maps of specific enhancer-promoter interactions^17^. We designed MCC experiments to assess promoter interaction profiles of five genes that showed enhanced response to LPS, DEX, and IFNG following IL-4 priming in a *Stat6*-dependent manner (enhanced LPS response - *Slc7a2, Ccl2* and *Edn1*; enhanced IFNG response – *Ciita*; enhanced DEX response - *Klf4*) (**Figure 2A and D**). MCC revealed remarkably specific IL-4-induced promoter-enhancer contacts that also exhibited stability after cytokine washout for each of these gene promoters. Specifically, we identified IL-4-induced promoter-enhancer contacts in the *Slc7a2* genomic locus with a −11kb *cis*-element, in the *Klf4* genomic locus with a −92kb *cis*-element, and in the *Ciita* genomic locus with a −27kb *cis*-element. Importantly, in the model of short-term transcriptional memory, these chromatin interactions were fully retained after IL-4 washout (**Figure 2D**). Closer inspection of the Memory loop anchor points revealed that these closely overlapped with OCRs defined by ATAC-seq, and either show IL-4-induced transient change in accessibility (*Slc7a2*: −11kb), or do not associate with noticeable changes in chromatin accessibility (*Klf4*: −92kb and *Ciita*: −27kb) (**Figure 2D**). Further, these anchor points in these instances were not CTCF binding sites, but showed IL-4-induced STAT6 binding, indicating that STAT6 is important for the formation of these chromatin loops. Strikingly, these same regions readily bound the main effector TFs in the presence of secondary stimulants (i.e., LPS-activated p65 - NF-kB complex; DEX-activated glucocorticoid receptor – GR; and IFNG-activated STAT1). Finally, we found that the expression profiles of *Slc7a2, Klf4*, and *Ciita* (from RNA-seq) showed IL-4-driven induction, which was greatly enhanced by the addition of the secondary stimulants in both the Signal integration and Short-term transcriptional memory settings in an IL-4/STAT6-dependent manner. We observed very similar results for two additional LPS-regulated genes, *Ccl2*, and *Edn1*; however, *Ccl2* exhibited multiple pre-formed, promoter-enhancer loops in resting MFs, many of which were further enhanced by the addition of IL-4, with one particular anchor point exhibiting greatly enhanced interaction frequency between the promoter of *Ccl2* and a CTCF-bound *cis*-element (**Figure S2C**). Together, MCC revealed remarkably specific IL-4-induced promoter-enhancer loops that are stable and readily bound by the effector TFs of secondary stimulants, reinforcing the notion that these new loop anchor points are required for integrating unrelated, or even opposing inflammatory signals and might have functional relevance in the development of short-term transcriptional memory.

### IL-4-driven chromatin conformation is required for signal integration and memory formation in the *F10* gene locus

We set out to perturb IL-4-induced loop anchor points that associate with inflammatory signal integration and short-term transcriptional memory formation. We decided to interrogate the *F7*/*F10* gene locus that exhibited stable IL-4-induced loops in the Hi-C data and are considered as alternative MF polarization marker genes (Figure 1F). We designed MCC experiments and targeted both genes’ promoters, which recapitulated the Hi-C observations and provided us with high-resolution promoter-mediated contact maps in the priming system, revealing that the promoters exhibit IL-4 induced stable looping – after washout – with not only its cognate enhancers (e.g., −15kb), but also with two CTCF-bound sites (located −26kb and +68kb from the TSS of *F10*) that most likely confine this gene regulatory unit and represent a topologically-associated domain (TAD) (**Figure 3A**).

**Figure 3.**
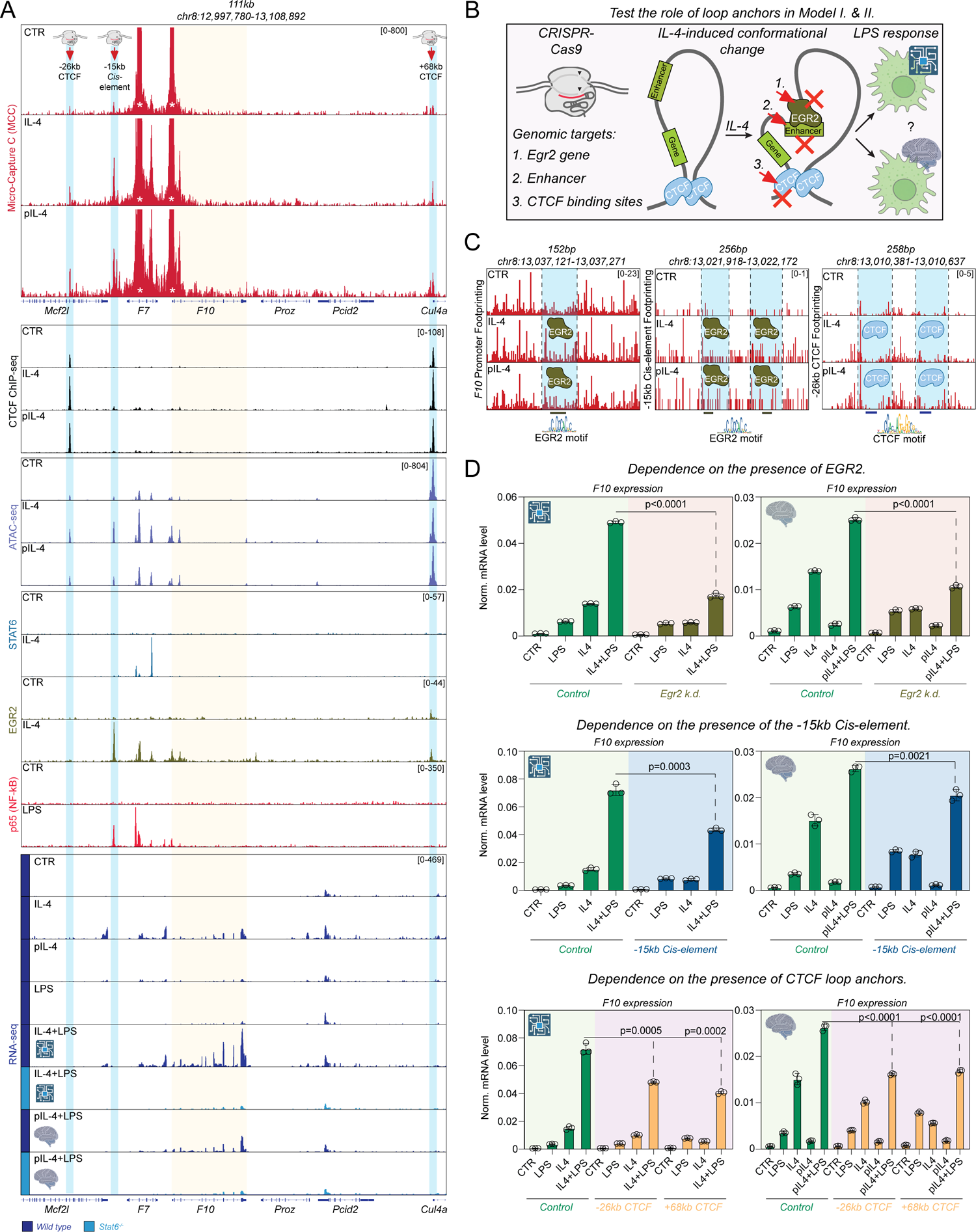
Memory loop anchors are functional, integrate opposing polarization signals, and confer short-term transcriptional memory. **(A)** Genome browser tracks showing different datasets in the *F10* gene locus that nominated three loop anchor points for CRISPR editing. **(B)** Schematics that present the hypothesis to be tested that states that the IL-4-induced stable loop anchors are functional and required for enhanced LPS response. **(C)** Base-pair resolution transcription factor footprinting by plotting the ligation junctions in the indicated *cis*-elements. **(D)** RT-qPCR measurements of *F10* gene expression in the signal integration (microchip icon) and short-term memory (brain icon) settings upon various CRISPR perturbations. Significant differences were determined by two tailed, unpaired t-test at p<0.05.

*Cis*-elements that are detected in this TAD exhibited strongly elevated ATAC-seq signal in the presence of IL-4 and retained their openness after cytokine washout. Among these *cis*-elements, the −15kb *cis*-element readily bound EGR2, an IL-4/STAT6 induced TF with long-term epigenetic effects in alternatively polarized MFs ^7, 30, 31^. Additionally, this *cis*-element also bound LPS-activated p65; however, the most upstream transcriptional regulator of IL-4-induced transcription, STAT6, was not detected at this site, but rather at a neighboring *cis*-element that is located in the intron of the *F7* gene (**Figure 3A**). This observation aligns with recent reports that STAT6 is an upstream regulator of EGR2’s expression, and the two factors’ cistromes (the sum of all binding sites in the genome) show a largely mutually exclusive genomic binding pattern (i.e., spatially and temporally separated cistromes) in the MF genome upon alternative MF activation^7, 31^. RNA-seq revealed that only *F10* exhibits signal integration and short-term memory formation specifically in the IL-4 – LPS setting, but *F7* is insensitive to LPS (**Figure 3A**). Therefore, we focused on the *F10* genomic locus and performed base-pair resolution chromatin conformation mapping with MCC^17^. To achieve near base-pair resolution analysis, we plotted the exact positions of ligation junctions between the gene promoter and the respective, interacting *cis*-elements, revealing potential TF footprints for EGR2 in the gene promoter and in the −15kb *cis*-element. We also uncovered two strong CTCF footprints in the −26kb CTCF-bound region (**Figure 3C**). These results nominated the −15kb *cis*-element, and the two CTCF sites located −26kb and +68kb from *F10* for CRISPR perturbation experiments in primary MFs (**Figure 3B**).

First, we designed single guide RNAs (sgRNA) to target the *Egr2* gene to assess the requirement of this TF; second, we designed sgRNAs to perturb the −15kb *cis*-element; and third, we designed sgRNAs to target the two CTCF sites in the models of Signal integration and Short-term transcriptional memory formation (**Figure 3B**). After optimization of electroporation conditions using the cell surface expression of CD11B (ITGAM) as a surrogate according to a recent report (**Figure S3A**)^32^; we first perturbed *Egr2* expression leading to a ∼45% reduction in IL-4-induced *Egr2* expression at the mRNA level (**Figure S3B**), which was followed by severely reduced signal integration and short-term transcriptional memory formation ability in MFs as measured by RT-qPCR of *F10* mRNA levels (**Figure 3D**). Similarly, when we perturbed the − 15kb *cis*-element, - that as a result of perturbation, lost its ability to respond to LPS as assessed by RT-qPCR measurements of enhancer RNA (eRNA) production (**Figure S3C)** - we found that this *cis*-element was also required in both treatment paradigms for enhanced transcriptional response to LPS, although its perturbation had a milder impact on *F10* gene expression than the loss of *Egr2*. Finally, we perturbed the two CTCF sites (−26kb and +68kb), which both led to reduced signal integration and short-term transcriptional memory formation after IL-4 exposure as measured by RT-qPCR of *F10* mRNA levels (**Figure 3D**). Collectively, these results indicate that IL-4-induced chromatin conformation and the anchor points of the new conformation are required for enhanced LPS-driven expression of *F10,* and the newly installed conformation exhibits stability and can provide transcriptional memory.

### Transcriptionally silent, IL-4-driven 3D chromatin conformation is required for enhanced endotoxin-induced *Il6* expression

Finally, we focused on *Il6*, a proinflammatory cytokine and a cardinal gene marker of innate immune memory formation with multiple roles in inflammatory responses, including being the key driver of cytokine storm^26, 33^. Importantly, *Il6* shows enhanced LPS response after IL-4 exposure in both the Signal integration and Short-term transcriptional memory settings; however, IL-4 does not impact the 1D epigenetic state of the *Il6* locus (Figure 1F), nor does it affect *Il6* expression (as opposed to *F10* in Figure 3). We performed MCC and found that the gene promoter contacts a *cis*-element in an IL-4-dependent manner that is located 64kb upstream from the gene, and this loop exhibits stability after IL-4 removal (**Figure 4A**), recapitulating the Hi-C results in Figure 1F. This *cis*-element exhibited weak enrichment for CTCF but showed strong ATAC-seq signal that was unaffected by IL-4; however, it readily bound STAT6 and EGR2 in response to IL-4. Additionally, in the presence of LPS, this *cis*-element binds p65; thus, during priming, IL-4 juxtaposes this element to the *Il6* promoter, primes the 3D chromatin conformation of the locus in the absence of any noticeable change in chromatin accessibility, histone modification, and gene expression (**Figure 4A**). Importantly, IL-4-activated *Stat6* is indispensable for enhanced LPS-driven *Il6* expression at the mRNA level as detected by RNA-seq (**Figure 4A**).

**Figure 4.**
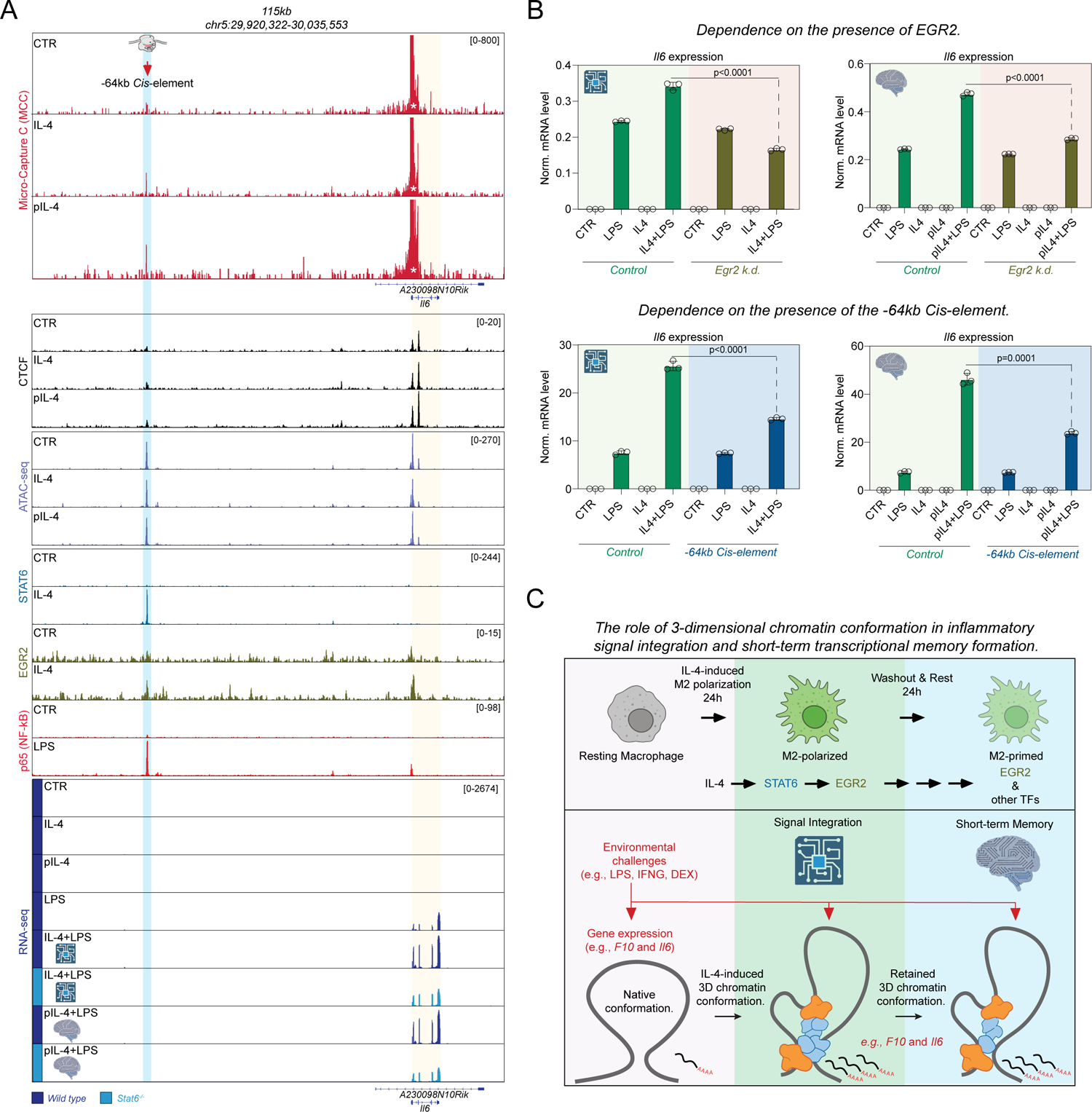
IL-4-induced memory loops can form in the absence of epigenetic remodeling and transcription. **(A)** Genome browser tracks showing different datasets in the *Il6* gene locus that nominated a loop anchor points for CRISPR editing. **(B)** RT-qPCR measurements of *Il6* gene expression in the signal integration (microchip icon) and short-term memory (brain icon) settings upon various CRISPR perturbations. Significant differences were determined by two tailed, unpaired t-test at p<0.05. **(C)** Schematics that summarize the main findings of the study. Briefly, IL-4 activated STAT6 can induced the expression of a secondary transcription factor, EGR2 in the transcription factor cascade that remodels the epigenetic and 3D genome conformation of thousands of genomic loci, some of which remains stable after the removal of IL-4. IL-4-induced chromatin loop anchor points are required for enhanced LPS response in the signal integration and short-term memory settings and provide elevated *F10* and *Il6* mRNA transcript levels.

We perturbed the expression of *Egr2* with CRISPR and found that *Egr2* was required for enhanced LPS response in *Il6* expression in both the Inflammatory signal integration and Short-term transcriptional memory settings as detected by RT-qPCR (**Figure 4B**). Finally, when we targeted the −64kb *cis*-element - which as a result of perturbation showed severely reduced LPS-driven eRNA levels (**Figure S3D**) - we observed reduced synergy between IL-4 and LPS in driving enhanced *Il6* expression in both the experimental systems of Signal integration and Short-term transcriptional memory formation (**Figure 4B**). To summarize, our results show that two unrelated, and in fact, opposing inflammatory signals (IL-4 and LPS) can collaborate solely through 3D chromatin conformation where the anchor points of the conformation are set by IL-4, and these anchor points are indispensable for enhanced LPS-triggered inflammatory response when the two signals are present at the same time, and can also provide short-term transcriptional memory (**Figure 4C**).

## Discussion

Here, we report critical functions of genome conformation in inflammatory signal integration and short-term transcriptional memory formation in primary MFs. We report that IL-4 can largely reorganize the 3D genome conformation of MFs, establish thousands of chromatin loops that are stable, even after the removal of the cytokine. We find that these Memory loops are enriched around IL-4-regulated genes, but also around genes that are not controlled by IL-4 at the transcriptional level; and surprisingly, we report that many of these gene loci do not exhibit any apparent 1D epigenetic remodeling. We show that hundreds of these genes with remodeled 3D chromatin conformation exhibit enhanced responsiveness to secondary stimulants (LPS, DEX, and IFNG). We used high-resolution MCC to pinpoint the anchor points of IL-4-driven loops that originate from gene promoters, and report that IL-4 juxtaposes LPS-, DEX-, and IFNG-activated *cis*-elements to their cognate gene promoters, a conformation that lasts for at least one day following the removal of IL-4. We perform CRISPR perturbation experiments to study the functional relevance of IL-4-induced memory loops in two independent gene loci, *F10*, where IL-4 can activate gene expression and remodels the epigenetic state of the locus during memory loop formation, and *Il6*, where IL-4 has no effect on the basal transcription of the gene, nor at the epigenetic state of the locus during memory loop formation. We show that both *F10* and *Il6* exhibit enhanced LPS-responsiveness after IL-4 priming when both IL-4 and LPS are present (Signal integration), and when IL-4 is removed and LPS stimulation is performed one day later. We find that the STAT6-activated TF, EGR2, is required for signal integration and short-term memory formation, and further, that EGR2 and LPS-activated p65 bind the anchor points of IL-4-established chromatin conformation and these genome architectural points are critical for signal integration and short-term transcriptional memory formation. Additionally, we report IL-4-driven Memory loops between the promoter of *F10* and CTCF-bound insulator-elements that are critical for signal integration and memory formation. Our results uncover fundamental new functions of the 3D genome conformation of specific gene loci in integrating unrelated environmental signals, and in establishing short-term memory formation, all of which can ultimately regulate MF plasticity.

Multiple studies have reported that IL-4 or other priming signals can reprogram the epigenetic landscape of MFs; however, in the absence of high-resolution genome conformation mapping, all of these studies were unable to assess the regulatory role of chromatin loops in the subsequent responses of MFs^3, 34, 35^. Since enhanced LPS-induced *Il6* expression following beta glucan (BG) priming is one of the markers of trained innate immunity, we suggest that a similar, 3D genome priming phenomenon can operate in the classical model of trained innate immunity as well; however, BG induces the expression level of *Il6*, whereas IL-4 does not. Further, we find that as a result of IL-4 priming, LPS can drive the gene expression programs of cholesterol metabolism and the mevalonate pathway - two related metabolic pathways that are critical for trained innate immunity via the production of metabolic intermediates^36^. Interestingly, IL-4-activated STAT6 turns on the expression of EGR2 and MITF TFs^7, 31^, similar to how BG training induces the expression of both TFs that associate with the rescue of LPS-tolerized human MFs from a suppressed state, not responsive endotoxin stimulation^35^. Importantly, in our system, the IL-4-STAT6-EGR2 cascade is required for enhanced LPS response, and we suggest that the activity of this TF cascade is a requirement for Memory loop formation, and that this mechanism can be analogous to trained innate immune memory formation.

Recent studies suggest that in some instances, active transcription may drive 3D chromatin conformational changes although the order of these events remains unresolved^37–40^. We find that IL-4 can trigger genome conformational changes without detectable changes in epigenetic state or transcription. In the *Il6* locus, we find that IL-4 creates a *de novo* chromatin loop that forms in the absence of any detectable epigenetic and transcriptional change and connects an LPS-activated p65-driven *cis*-element to the *Il6* promoter - an anchor point that is indispensable for proper signal integration and memory formation, leading to enhanced LPS response. These results indicate that cellular memory formation can be encoded in a specific 3D genome configuration – without the necessity of a transcriptional response – where the spatial constellation of epigenetic modifiers/modifications and *trans*-elements are required for enhanced transcriptional responses.

Overall, our work suggests that MF 3D genome conformation is dynamically regulated during polarization, and that this conformation has functional implications in integrating inflammatory signals and in the establishment of short-term transcriptional memory. These findings will facilitate future studies to better understand the broader roles of genome conformation in gene expression regulation and in cellular memory formation. We envision that one aspect of cellular memory formation can be encrypted in specific 3D folding of genomic regions as it has been suggested by computational modeling in the model of cell identity maintenance during and after cell division^54^. Future work will assess the role of 3D chromatin configuration in additional cell types of the immune system, particularly in long-lived memory B and T cells that provide long-term immunity against pathogens and are crucial for vaccination-induced immunity.

## Acknowledgements

We thank the members of the Satpathy lab for stimulating discussions. We thank James O. J. Davies for discussions and for sharing the Micro-Capture C protocol. This work was supported by the National Institutes of Health (NIH) U01CA260852 (A.T.S.), a Career Award for Medical Scientists from the Burroughs Wellcome Fund (A.T.S.), a Lloyd J. Old STAR Award from the Cancer Research Institute (A.T.S.), an ASH Scholar Award from the American Society of Hematology (A.T.S.), and the Parker Institute for Cancer Immunotherapy (H.Y.C., and A.T.S). H.Y.C. is an investigator of the Howard Hughes Medical Institute. The sequencing data was generated with instrumentation purchased with NIH funds: S10OD018220 and 1S10OD021763.

## Author contributions

B.D. and A.T.S conceptualized the study. B.D. and A.T.S. wrote and edited the manuscript and all authors reviewed and provided comments on the manuscript. B.D., K.S., W.Z., and Z.M. performed experiments. A.Y.C. set up computational pipelines and analyzed the data. K.E.Y. and C.A.L. performed computational analyses. B.D., H.Y.C. and A.T.S. guided experiments and data analysis.

## Declaration of interests

A.T.S. is a founder of Immunai, Cartography Biosciences, and Prox Biosciences, an advisor to Zafrens, Pallando Therapeutics, and Wing Venture Capital, and receives research funding from Merck Research Laboratories. H.Y.C. is a co-founder of Accent Therapeutics, Boundless Bio, Cartography Biosciences, Orbital Therapeutics, and an advisor of 10x Genomics, Arsenal Biosciences, Chroma Medicine, Exai Bio, and Spring Discovery. B.D. is an employee of Genentech. K.S. is an employee of Cartography Biosciences. K.E.Y. is a consultant for Cartography Biosciences. C.A.L. is a consultant of Cartography Biosciences.

## Data availability

Reviewer access for sequencing data is available under GEO accession: GSE247946. Reviewer token to access data: otwbkegqrtyxbgz

## Supplementary Figure Legends

**Figure S1. Epigenetic characterization of macrophage priming by IL-4. Related to** Figure 1**. (A)** Heatmap representation of IL-4-repressed chromatin loops determined by Hi-C. **(B)** Heatmap representation of IL-4-repressed mRNA transcripts (left). Small heatmaps represent the results of Ingenuity pathway analysis for each RNA-seq gene cluster, and the top 3 biological terms are shown (right). Extended data figure 2. **(C)** Heatmap representation of genomic regions with differential histone enrichment profiles and ATAC-seq profiles in the indicated conditions (IL-4 mediated priming model). **(D)** Venn diagrams depict the IL-4-induced and -reduced histone modification landscape and identifies the stable effects of IL-4 that is related to memory formation. Percentage wise distribution of the genomic regions across the CTR vs. IL-4 and CTR vs. pIL-4, and their overlap is plotted. **(E)** Scheme of annotation of distant IL-4-induced open chromatin regions to differentially expressed genes in a +/-100kb genomic window around their transcription start sites. **(F)** Histograms show the epigenetic state of IL-4-induced differentially accessible (top row) and non-differentially (bottom row) accessible cis-elements in the proximity (+/-100kb around the TSS) of not-expressed genes that have memory loops overlapping with their gene bodies from Fig. 1e (red section of the Venn diagram).

**Figure S2. Inflammatory signal integration is widespread. Related to** Figure 2**. (A)** Heatmap representation of differentially expressed genes (DEGs) detected by RNA-seq (left). Small heatmaps represent the results of Ingenuity pathway analysis for each RNA-seq gene cluster, and the top 3 biological terms are shown (middle). Permutation test results that assess the enrichment of Memory loops in the different groups of DEGs in the IL-4 – IFNG treatment paradigm. **(B)** Heatmap representation of differentially expressed genes (negatively affected by IL-4) in the indicated treatment paradigms and models. **(C)** Genome browser tracks showing genomic datasets in the *Ccl2* and *Edn1* gene loci.

**Figure S3. Transcriptional memory formation and signal integration is linked to specific memory loop anchor points. (A)** FACS data of CD11b expression on the cell surface of macrophages in the presence of a control guide RNA (ROSA-targeting) and Cd11b-targeting guide during a CRISPR optimization experiment. **(B)** RT-qPCR measurements of *Egr2* mRNA expression after control (ROSA-targeting) and *Egr2*-targeting guide and Cas9 electroporation. Significant differences were determined by two tailed, unpaired t-test at p<0.05. **(C)** Genome browser snapshot of p65 ChIP-seq and GRO-seq datasets that guided the design of enhancer RNA-specific primers to detect the effects of guide RNAs in CRISPR experiments that targeted these cis-elements. RT-qPCR measurements of *F10 - 15kb* eRNA expression after control (ROSA-targeting) and *cis-element*-targeting guide and Cas9 electroporation. **(D)** Same as panel C, but for the *Il6-64kb* cis-element. Significant differences were determined by two tailed, unpaired t-test at p<0.05.

## Methods

### Mice and macrophage differentiation

Wild type, 2-3 months old female C57/Bl6 mice were purchased from Jackson laboratories. Female *Stat6^-/-^* mice were purchased from Jackson Laboratories. Mice were sacrificed and bone marrow was isolated form the tibiae and femora of the animals. Red blood cell lysis was carried out and cells were plated in differentiation media containing 10% FBS, Dulbecco’s Modified Eagle’s Medium (DMEM) and 20ng/ml mouse M-CSF (Peprotech). On the third day of differentiation, media was replaced with fresh differentiation media. Cytokine treatments, sorting procedures were carried out on the 6^th^ day of differentiation.

### Treatment conditions

On the 6^th^ day of bone marrow-derived macrophage differentiation, cells were either left untreated, or were exposed to mouse IL-4 (20ng/ml; Peprotech) for 24-hours. For testing signal integration and short-term memory formation, following the 24-hour IL-4 treatment, cells were either left in IL-4 and immediately treated for 3-hours with the secondary stimulants (LPS – 100ng/ml, DEX – 100nM, IFNG – 20ng/ml), or IL-4 washout was performed (3-times with PBS after the removal of IL-4 containing media), cells were further incubated for 24-hours in differentiation media followed by secondary stimulant treatment for 3-hours, respectively.

### CRISPR-based gene and *cis*-element engineering

On the 5^th^ day of bone marrow-derived macrophage differentiation, cells were scraped in differentiation media, spun, and resuspended and counted in PBS. Before the cells were prepared and counted for electroporation, Cas9-Ribonucleoprotein (RNP) complexes were prepared as follows: crRNAs and tracrRNA were resuspended to 100uM in resuspension buffer (IDT). For each electroporation reaction, 2ul crRNA was mixed with 2ul tracrRNA for duplex formation. Duplex formation was carried out as follows: crRNA/tracrRNA complexes were heated to 95C for 5-minutes followed by cooling to 20C at 0.5C per second, and were stored on ice until it was mixed with recombinant Cas9 at 3:1 molar ratio. RNP complexes were made in the following reaction while incubating at room temperature for 20-minutes: 4ul RNA duplex, 1.33ul 10mg/ml recombinant Cas9, and 1ul H_2_O. After the Cas9-RNP complexes were made, macrophages were scraped and counted to achieve 0.5×10^6^ cells were prepared in 20ul P3 buffer for electroporation (LONZA - 4D Nucleofactor). To each 20ul macrophage suspension in P3 buffer, 5ul of Cas9-RNP complex solution was added and mixed 5-times before 24ul was transferred to an electroporator well (pipetted two times in 12ul volumes). Electroporation was performed with the CM-137 setting as described previously^32^. Immediately after electroporation, 180ul differentiation media (including 20ng/ml M-CSF) that was pre-warmed to 37C was added to each well and cells were resuspended and plated to 12 well plates in a total of 400ul macrophage differentiation media. Treatments were performed on the next day as described above in the Treatment conditions section that was followed by incubation and either flow cytometry or RNA isolation and real-time quantitative PCR measurements. Guide RNA sequences are available from Table S8.

### RNA isolation

RNA was isolated from 2×10^5^ bone marrow-derived macrophages (BMDMs) by standard, TRIzol-based RNA precipitation method as follows. Cells were resuspended in 1ml TRIzol (Ambion). Chloroform was added (200ul) to this lysate and extensively vortexed to achieve a homogenous mixture; then, it was incubated for 3-minutes at room temperature before centrifugation at 14,000 rcf (relative centrifugal force) at 4C for 15-minutes. Aqueous layer was collected from the top and transferred into a new tube (∼550ul), 1ul GylcoBlue (Ambion) was added, and the RNA was precipitated with equal volume of 2-propanol for 20-minutes at room temperature. RNA precipitates were centrifuged at 16,000 rcf for 15-minutes at 4C and supernatant was carefully discarded without disturbing the GlycoBlue-stained, blue, RNA pellet. RNA pellet was washed with 1ml 75% EtOH and after the wash, dissolved in 30ul nuclease-free water. RNA concentration was determined by nanodrop, and RNA quality was determined by Agilent Bioanalyzer.

### Real-Time quantitative PCR (RT-qPCR)

RNA was isolated with Trizol reagent (Ambion). RNA was reverse transcribed with High-Capacity cDNA Reverse Transcription Kit (Applied Biosystems) according to the manufacturer’s instructions. Transcript quantification was performed by qPCR reactions using SYBR green master mix (BioRad). Transcript levels were normalized to *Ppia*. Primer sequences are available from Table S8.

### Fluorescence-activated cell sorting (FACS)

Macrophages (∼0.5 x 10^6^) were treated with Fc-block (1:100) for 15-minutes on ice, and then, stained with anti-ITGAM (rat monoclonal BV-510-conjugated, BioLegend) in a 1:200 dilution in FACS buffer (0.1% BSA, 2mM EDTA, 5% FBS in PBS) for 20-minutes on ice. Cells were spun and resuspended in PBS with LIVE/DEAD Fixable Aqua cell stain and incubated for 10-minutes on ice followed by FACS analysis.

### RNA-seq

Approximately, 50ng RNA was reverse-transcribed to complementary DNA (cDNA) and second strand was synthesized by the Ovation RNA-seq System V2 (Tecan) according to the manufacturer’s recommendations. Double-stranded DNA was subjected to isothermal amplification and was purified with Ampure XP beads. DNA was quantified by Qubit, and 80 ng DNA was used for sequencing library construction with the Ovation Ultralow Library System V2 (Tecan) using 8 PCR cycles according to the manufacturer’s recommendations. Libraries were sequenced with Illumina Novaseq 6000, using paired-end 75bp read configuration.

### ChIP-seq

ChIP-seq was performed as previously described with the following modifications^41^. Bone marrow-derived macrophages (5 x 10^6^) were double crosslinked by 50mM DSG (disuccinimidyl glutarate, #C1104 - ProteoChem) for 30-minutes followed by 10-minutes of 1% formaldehyde. Formaldehyde was quenched by the addition of glycine. Nuclei were isolated with ChIP lysis buffer (1% Triton x-100, 0.1% SDS, 150 mM NaCl, 1mM EDTA, and 20 mM Tris, pH 8.0). Nuclei were sheared with Covaris sonicator using the following setup: Fill level – 10, Duty Cycle – 5, PIP – 140, Cycles/Burst – 200, Time – 4 minutes). Sheared chromatin was immunoprecipitated overnight with the following antibodies: H3K4me3 (Diagenode – C1540003-50), H3K27ac (Abcam – ab4729), H3K4me1 (Diagenode – C15410194), and CTCF (Abcam – ab70303). Antibody chromatin complexes were pulled down with Protein A magnetic beads and washed once in IP wash buffer I. (1% Triton, 0.1% SDS, 150 mM NaCl, 1 mM EDTA, 20 mM Tris, pH 8.0, and 0.1% NaDOC), twice in IP wash buffer II. (1% Triton, 0.1% SDS, 500 mM NaCl, 1 mM EDTA, 20 mM Tris, pH 8.0, and 0.1% NaDOC), once in IP wash buffer III. (0.25 M LiCl, 0.5% NP-40, 1mM EDTA, 20 mM Tris, pH 8.0, 0.5% NaDOC) and once in TE buffer (10 mM EDTA and 200 mM Tris, pH 8.0). DNA was eluted from the beads by vigorous shaking for 20 minutes in elution buffer (100mM NaHCO_3_, 1% SDS). DNA was decrosslinked overnight at 65C and purified with MinElute PCR purification kit (Qiagen). DNA was quantified by Qubit and 10 ng DNA was used for sequencing library construction with the Ovation Ultralow Library System V2 (Tecan) using 12 PCR cycles according to the manufacturer’s recommendations. Libraries were sequenced with Illumina Nextseq 550, using paired-end 75bp read configuration.

### ATAC-seq

ATAC-seq was performed by using 10^5^ bone marrow-derived macrophages (BMDMs) from each condition. Nuclei were isolated with ATAC Lysis Buffer (10mM Tris-HCl pH7.4, 10mM NaCl, 3mM MgCl_2_, 0.1% IGEPAL). Nuclei from BMDMs were subjected to tagmentation using Nextera DNA Library Preparation Kit (Illumina). After tagmentation DNA was purified with MinElute PCR Purification Kit (Qiagen). Tagmented DNA was then amplified with Phusion high-fidelity PCR master mix (NEB) using 14 PCR cycles. Amplified libraries were purified again with MinElute PCR Purification Kit. Fragment distribution of libraries was assessed with Agilent Bioanalyzer and libraries were sequenced on a Novaseq 6000 platform with 150bp paired-end sequencing.

### Hi-C

Hi-C was performed by following the reported HiChIP protocol and omitting the protein immunoprecipitation step^42^. Briefly, 10^6^ bone marrow-derived macrophages were crosslinked with 1% formaldehyde for 10-minutes. Formaldehyde was quenched by the addition of glycine. Nuclei were isolated in lysis buffer (1% Triton x-100, 0.1% SDS, 150 mM NaCl, 1mM EDTA, and 20 mM Tris, pH 8.0) by rotating the cells for 1-hour at 4C. Nuclear pellet was resuspended in 0.5% SDS solution and incubated at 62C for 10 minutes. SDS was quenched by the addition of 10% Triton-X. Chromatin was digested with 8U of MboI enzyme overnight at 37C. Biotin fill-in was performed for 1-hour at 37C, followed by proximity ligation for 6-hours at room temperature. After proximity ligation, nuclei were subjected to Proteinase K digestion and decrosslinking overnight at 68C. DNA was column purified (Minelute, Qiagen), and sonicated to a size distribution of 200-500bp. DNA was subjected to streptavidin bead binding to pull-down biotinylated DNA that represents ligation junctions. Tn5 was used to create libraries by on bead tagmentation with sequencing adapters, followed by PCR. Libraries were quantified by Qubit and size distribution was assessed with Bioanalyzer. Libraries were sequenced on NovaSeq 6000 with paired-end 150bp configuration.

### Micro-Capture C

Micro-Capture C (MCC) was performed as previously described with the following differences^17^. Macrophages were scraped and crosslinked in the presence of 1% formaldehyde (in PBS) for 10-minutes in suspension followed by the addition of 1.5ml Glycine (1M) and further incubation for 5-minutes at room temperature. Cells were washed in ice-cold PBS, counted, and 3×10^6^ aliquots were prepared for nuclei isolation with the following MCC lysis buffer: 10mM Tris-HCl pH 7.5, 10mM NaCl, 3mM MgCl2, 0.1% Twee-20, 0.1% Nonidet P40 Substitute, 0.01% Digitonin and 1% BSA in nuclease-free H_2_O. Each cell aliquot of 3×10^6^ were resuspended in 500ul MCC lysis buffer and incubated on ice for 5-minutes. Cells were spun down, washed in ice-cold PBS, and washed in nuclease-free H2O before proceeding to MNase digestion with 1-2U of MNase for 3×10^6^ cells that went on for 1-hour at 37C while shaking in a thermomixer (Eppendorf) at 550rcf. MNase was stopped by the addition of EGTA. At this point, nuclei were split into two tubes, and end-repair and proximity ligation were performed overnight. On the next day, nuclei were spun and subjected to proteinase K digestion at 65C overnight. DNA was column purified (Minelute, Qiagen) and eluted in 130ul H2O for sonication with Covaris E220 model using the following settings: Fill Level – 10, Duty Cycle – 5, PIP – 140, Cycles/Bursts – 200; sonicated for 4-minutes. DNA was cleaned up with Minelute, quantified and subjected to library preparation with Ovation Ultralow V2 Library Systems (Tecan) using 100ng DNA and 8 PCR cycles. Amplified libraries were subjected to hybridization with biotinylated oligonucleotide pools that were designed by the HyperDesign Team (Roche) and synthesized by Roche. All hybridization reagents were purchased from Roche as part of the KAPA Hyper Prep kit, and hybridization, pull-down, and washes were performed per manufacturer’s instructions. Bead-bound libraries were amplified with the PCR reagents from the Ovation Ultralow Library V2 Kit (Tecan). Libraries were quantified and sequenced on a Novaseq 6000 with 2×150bp configuration.

### Micro-Capture C analysis

Micro-Capture C analysis was performed as described previously^17^. Briefly, adaptor sequences were removed using Trim Galore. Since the majority of the fragments were sonicated to 200bp, 2×150bp sequencing was performed, and the chimeric fragments were reconstructed into single sequences using FLASH. These reads were then mapped to a specific DNA sequence around the oligonucleotide with the help of the BLAT aligner, using an 800-bp region centered on the oligonucleotide as a reference. A second script, <u>MCCsplitter.pl</u>, was employed to split the reads based on their BLAT mapping. Reads that didn’t align with any capture oligonucleotide sites were discarded. The segmented reads were subsequently mapped to the genome using Bowtie2. Another script, <u>MCCanalyser.pl</u>, was used to further analyze the data, which is to remove PCR duplicates and to filter out regions with mapping issues. Ligation junctions were identified, and JASPAR was utilized to detect transcription-factor-binding sites at protected DNA regions that exhibited less frequent ligation points. Genome browser track signals in figures represent an average derived from at least six distinct replicates with different MNase concentrations.

### RNA-seq analysis

The sequencing data was analyzed using the nf-core RNA-seq pipeline (version 3.9) with default parameters (https://nf-co.re/rnaseq). Initially, FastQC was used for quality control of the fastq files, followed by read trimming via Trim Galore. These processed fastq files were then aligned to the mm10 mouse genome with STAR aligner. Subsequently, Salmon generated a gene-by-sample count matrix for further analysis. After performing PCA on variance-stabilized read counts, batch effects from donors were addressed using the removeBatchEffect function from the limma package^43^. DESeq2 package was used for the analysis of differentially expressed genes according to the following criteria: absolute log_2_ fold change of >=0.5 and FDR<0.05^44^. For visualization, a heatmap was constructed by aggregating differential genes of different treatment condition against control group and standardizing expressions. Genes were clustered using k-means clustering, considering Pearson correlation between rows. QIAGEN Ingenuity Pathway Analysis and clusterProfiler (version 4.8.3) were used for pathway analysis of genes in each cluster.

### ChIP-seq analysis

Sequencing adapters were trimmed using Trimmomatic and the reads were aligned to the mm10 reference genome using Bowtie2 with the --very-sensitive option^45^. The BAM files obtained were sorted based on their genomic coordinates using Samtools to eliminate PCR duplicates. After alignment and sorting, PCR duplicates were identified and removed using Picard. The BAM files were then converted to BED files using bamToBed and intersected with the mm10 blacklist. To normalize the output bedgraph files, genomeCoverageBed from the Bedtools package was used, and the resulting files were converted to bigwig format using bedGraphToBigWig from the UCSC tools. Peaks were identified from the BED file using MACS2 with a false discovery threshold of 0.05 (-q 0.05)^46^. To create a non-overlapping consensus peak set, overlapping peaks with lower significance across all samples were iteratively eliminated. By counting the overlaps of reads in each sample with the consensus peak set, a peak-by-sample matrix was generated. Differential peak analysis was then performed on the peak-by-sample count matrix, applying an absolute log2 fold change threshold of 0.5 and a p-value threshold of 0.05.

### Hi-C analysis

we utilized the nf-core/hic (2.0.0) pipeline for Chromosome Conformation Capture (Hi-C) data analysis, built on the Nextflow framework with Docker/Singularity containers ensuring reproducibility. Initial read quality was assessed using FastQC^47^. HiC-Pro was used for the core processing, including mapping with bowtie2, interaction product detection, and contact map generation^48^. Genome-wide contact maps were created and normalized using cooler^49^ and JuiceBox^50^. Data export, quality assessments, chromosomal compartments, and loop calling was conducted using Homer^51^. A final quality report was generated with MultiQC. The loops with 10k resolution were imported with hictoolsr (v 1.1.2) and HiC matrix counts were extract from .hic files. Differential loop heatmap constructed by aggregating differential genes (log2 fold change > 0.5 and p-value < 0.1) of different treatment condition against control group and standardizing counts.

### Loop anchor gene enrichment permutation test

Loop anchors were extended for 10 kb on both sides, and any genes that overlap with at least one of the extended anchors of interacting loops were considered regulated by the interacting loops. Permutation tests were performed with the permTest function in the regioneR package^52^. Briefly, we calculated the number of overlaps between the targeted genes and the loop anchors; then we selected 5,000 randomized genomic sets of regions of the same size and number using the randomizeRegions function in genomic to establish the background distribution and perform the statistical tests.

### Integrative analysis of RNA-, ATAC-, and ChIP-seq

Chromatin accessibility and ChIP binding signal histograms and heatmaps around the TSS of differentially expressed genes were generated using featureAlignedDistribution from ChIPseeker R package^53^. A 3 kb window was used to aggregate ATAC and ChIP track signals at differentially accessible sites around the upstream and downstream 100kb of the TSS.

### Statistical Analyses

Statistical analyses were performed in R or GraphPad Prism. qPCR measurements were presented as means +/- SD and three biological replicates were performed. The exact replicate numbers are indicated in the figure legends for each experiment. On the bar graphs, significant changes were determined by two tailed, unpaired t-test at p<0.05. Differential loops from Hi-C were called by p<0.1, Log_2_ FC≥0.5. Differential chromatin accessibility analyses across cell clusters were performed with the following parameters: FDR≤0.01, Log_2_ FC≥1, unless specified otherwise. Differential gene expression analyses were performed with the following parameters: FDR≤0.01, Log_2_ FC≥1. Statistical parameters are reported in the figure legends and in the results section.

