## Supplementary figures and images for "Regulation of immune signal integration and memory by inflammation-induced chromosome conformation"

### Supplemental Figures

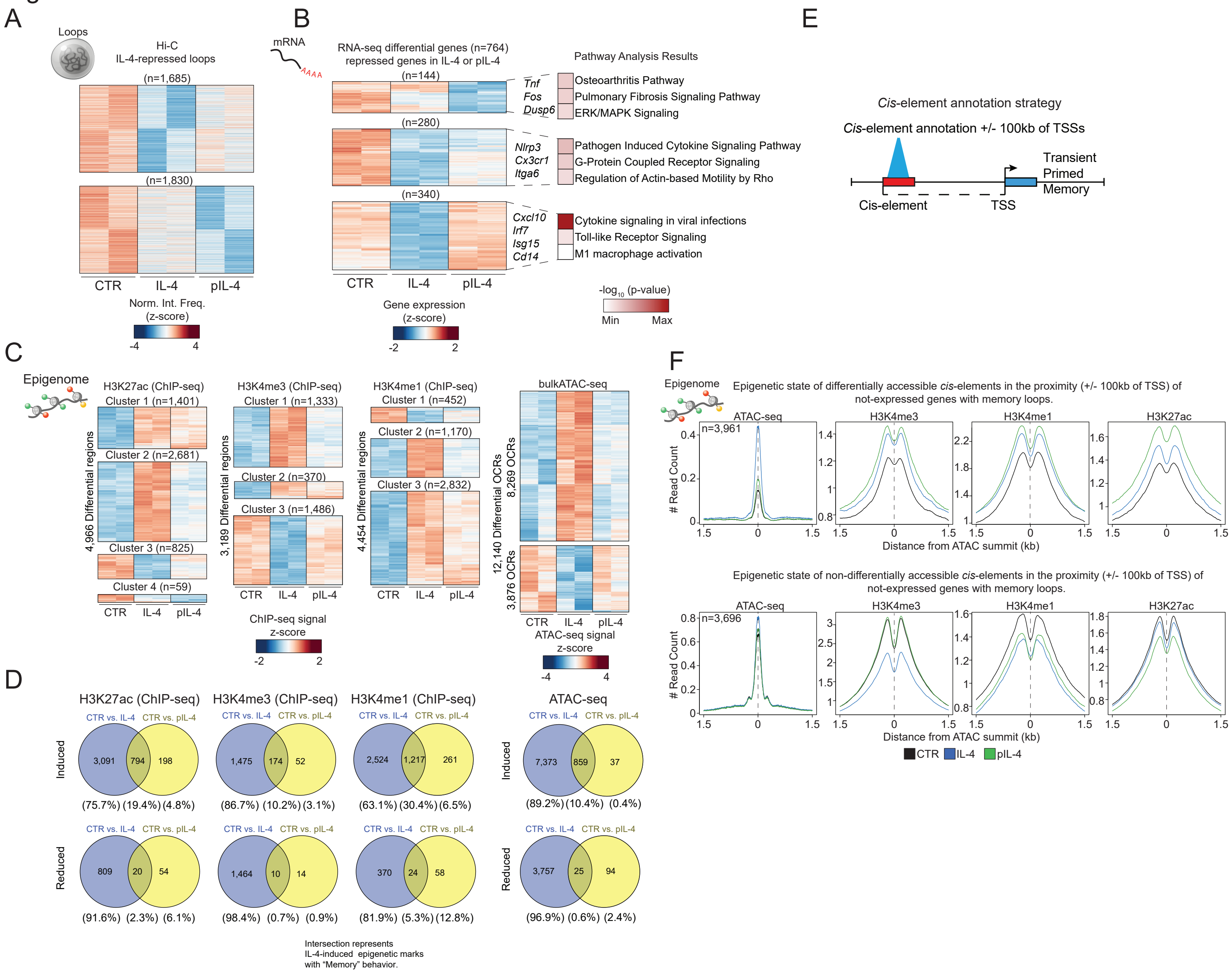

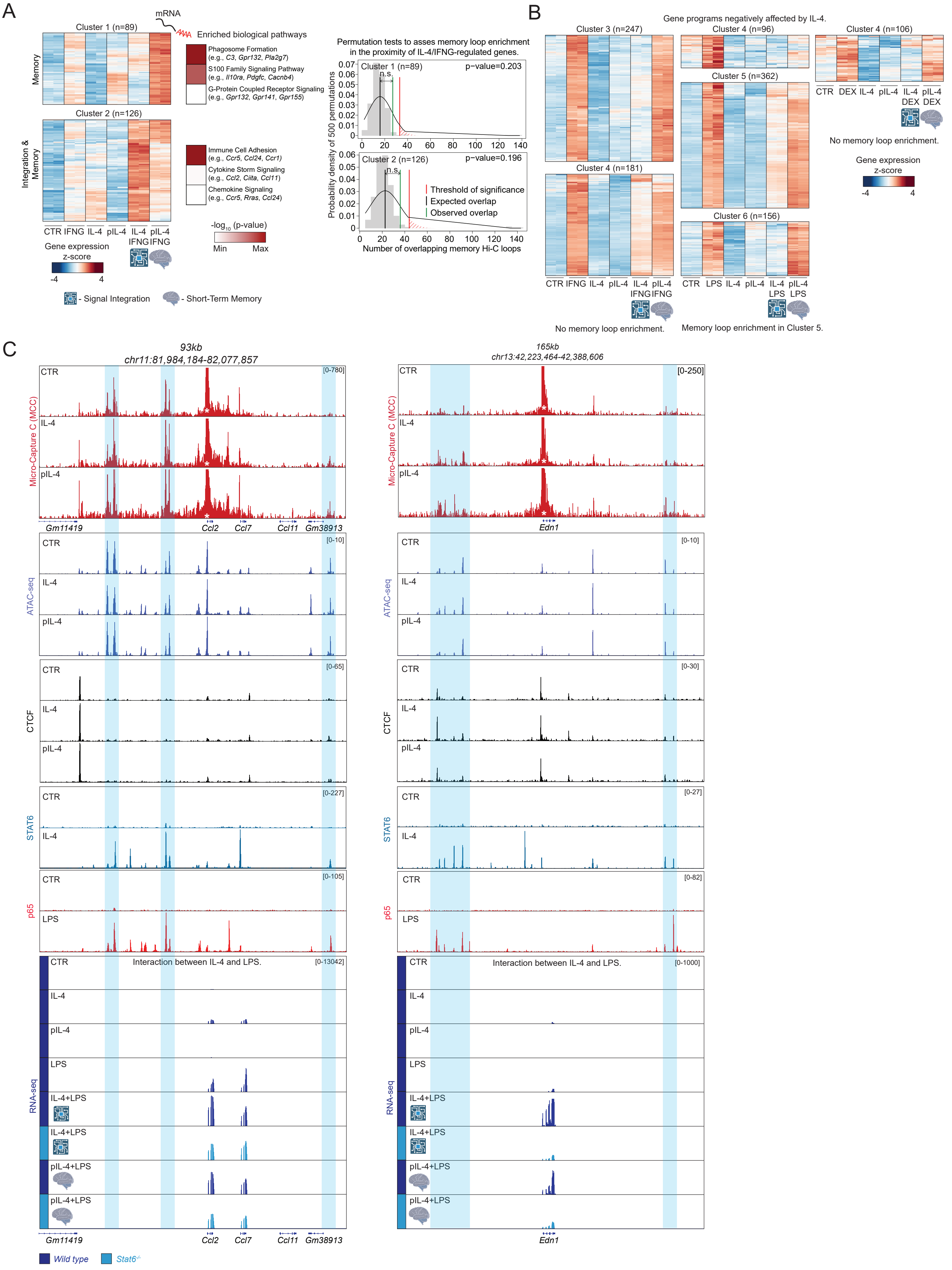

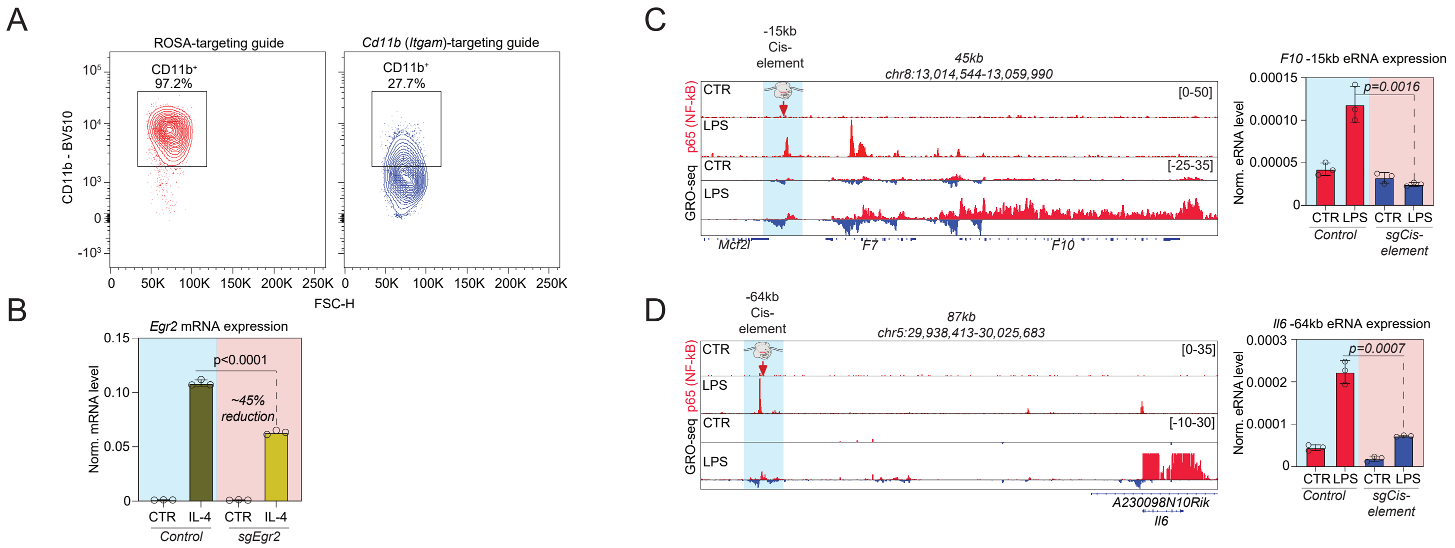
